# Black-necked spitting cobra (*Naja nigricollis)* phospholipases A_2_ cause *Trypanosoma brucei* death by blocking endocytosis through the flagellar pocket

**DOI:** 10.1101/2021.11.01.466813

**Authors:** Andrea Martos-Esteban, Olivia J. S. Macleod, Isabella Maudlin, Konstantinos Kalogeropoulos, Jonas A. Jürgensen, Mark Carrington, Andreas H. Laustsen

## Abstract

African trypanosomes, such as *Trypanosoma brucei,* are flagellated protozoa which proliferate in mammals and cause a variety of diseases in people and animals. In a mammalian host, the external face of the African trypanosome plasma membrane is covered by a densely packed coat formed of variant surface glycoprotein (VSG), which counteracts the host adaptive immune response by antigenic variation. The VSG is attached to the external face of the plasma membrane by covalent attachment of the C-terminus to a glycosylphosphatidylinositol. As the trypanosome grows, newly synthesised VSG is added to the plasma membrane by vesicle fusion to the flagellar pocket, the sole location of exo- and endocytosis. Snake venoms contain dozens of components including proteases and phospholipases. Here, we investigated the effect of *Naja nigricollis* on *T. brucei* with the aim of describing the response of the trypanosome to hydrolytic attack on the VSG. We found no evidence for VGS hydrolysis however *N. nigricollis* venom caused: (i) an enlargement of the flagellar pocket, (ii) the Rab11 positive endosomal compartments to adopt an abnormal dispersed localisation, and (iii) a cell cycle arrest prior to cytokinesis. A single protein family, the phospholipases A_2_s present in *N. nigricollis* venom, was necessary and sufficient for the effects. This study provides new molecular insight into *T. brucei* biology and possibly describes mechanisms that could be exploited for *T. brucei* targeting.

## Introduction

African trypanosomes are protozoan pathogens that have evolved the capacity to evade the mammalian innate and adaptive immune systems, and infection causes a variety of diseases in people and livestock^1^. In a mammalian host, the external face of *Trypanosome brucei* plasma membrane is covered by a densely packed coat formed by variant surface glycoprotein (VSG)^2^. VSG is at the centre of a population survival strategy based on antigenic variation, enabling the infection to avoid clearance by the host adaptive immune response (Fig. 1 A-I). The VSG is also central to individual cell survival strategy: Once an immunoglobulin (Ig) binds a VSG, the complex (VSG-Ig) protrudes above the surface coat and is subject to hydrodynamic forces, imposed by the swimming motion of the cell, causing the VSG-Ig to move towards the cell posterior pole and the flagellar pocket where it is endocytosed^3,4^. The flagellar pocket is an invagination within the plasma membrane^5^, from where the flagellum exits the cell body and is the sole site of endo- and exocytosis^3^ (Fig. 1 A-II). The rate of endocytosis is particularly high in trypanosomes^6,7^, and is all clathrin-mediated^8^. After endocytosis, Ig is trafficked through the internal endocytic pathway to the lysosome. This movement of cargo occurs through compartments that are highly polarised along an axis from the flagellar pocket to the lysosome, which lies close to the nucleus (Fig. 1 A-III)^9,10^.

**Figure 1.**
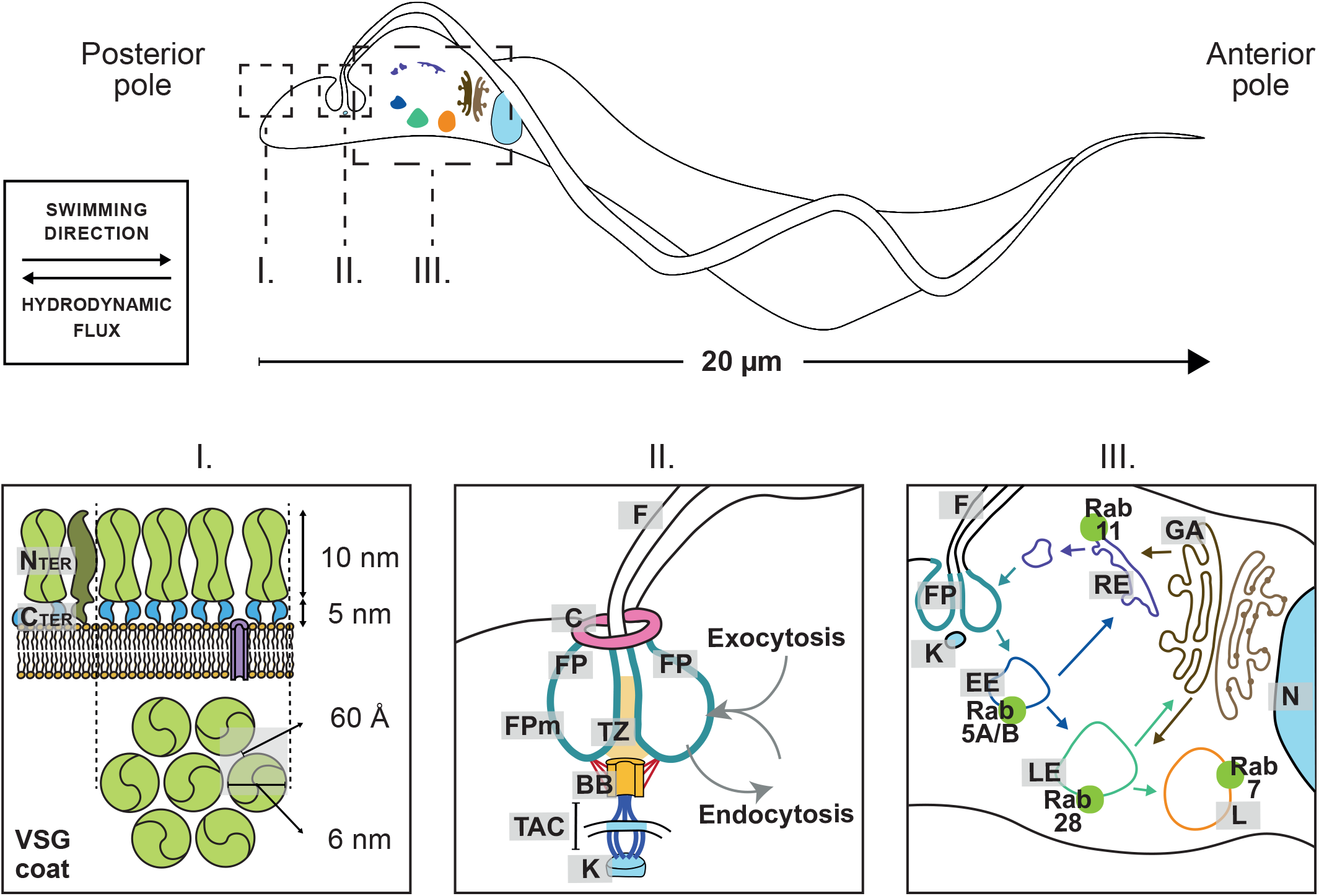
*T. brucei* morphology: I. VSG coat structure. The plasma membrane is covered by a densely packed VSG coat. VSG constitutes over 95% of the surface coat components^2^. **II. Flagellar pocket structure**^5^. The flagellar pocket (**FP**) is the sole site of endo- and exocytosis. This invagination is structured by a highly specialised membrane (**FPm**) penetrated by the flagellum (**F**) and stabilised by the flagellar pocket collar (**C**). The flagellum is attached to the cell body by the basal body (BB), which is connected by the tripartite attachment zone to the kinetoplast (**K**), the kinetoplastids’ unique organelle. **III. Endocytosis and exocytosis pathways in *T. brucei***^2,3^. Cargo is endocytosed in the flagellar pocket. The invaginated vesicles fuse to the Rab5A/B+ early endosome (**EE**). Then, cargo can be sent for recycling or for modification/degradation. Via the recycling pathway, cargo goes to the Rab11+ recycling endosome, and finally is exocytosed through the flagellar pocket. Should the cargo undergo modification/degradation, it is sent to the Rab28+ late endosome (**LE**). There, cargo is sent either for degradation via Rab7+ lysosome (**L**) or for modification via the Golgi apparatus.

Snake and bee toxins have been used as tools for the study of trypanosomatid-related diseases. In particular, several studies have shown the effect of various phospholipases in trypanosomatids. Phospholipase A_2_ (PLA_2_) present in *Apis mellifera* is lethal to *T. brucei*^11^; PLA_2_ BnSP-7 from *Bothrops pauloensis*^12^ and Asp49- and Lys49-PLA_2_s from *Bothrops matogrossensis*^13^ interfere with proliferation and infectivity of *Leishmania amazonensis*. In general, PLA_2_s from a range of venoms have a sequence identity of 40% to 99% and a conserved structure, with a four alpha-helix core, where each helix is joined by a loop or a sheet^14^. Despite sequence and structural homology, different subtypes of PLA_2_s have shown to play diverse functions and mechanisms of action^15,16^. However, the molecular pathways triggering trypanocidal effects and phenotypes caused by PLA_2_s have not been investigated.

*Naja nigricollis* (the black-necked spitting cobra) is one of the medically most important snakes found across West, Central, and East Africa^17^. It is known for having a venom rich in PLA_2_s and cytotoxic three-finger toxins that combined cause severe cytotoxicity, manifesting as local tissue damage and necrosis in mammalian victims. Here, we investigate the effects of *N. nigricollis* venom (NNV) on *T. brucei* at the mechanistic and molecular level. We show that NNV is trypanocidal and identify the lethal components to be PLA_2_s, which comprise ~22% of total venom protein^17^. We show that the PLA_2_s present in NNV are the first external effector causing massive flagellar pocket enlargement, the ‘Big Eye’ phenotype^8^. This phenotype was first described in trypanosomes in which clathrin light chain^8^ or actin^18^ had been depleted, and enlargement of the flagellar pocket occurs as membrane is still added by exocytosis, but none removed by endocytosis, eventually resulting in cell death.

## Results

### *N. nigricollis* venom causes *T. brucei* cell death

To assay for potential effects on the growth rate and survival, cultures of *T. brucei* were incubated with NNV over 24 hours (Fig. 2 A). By comparison to a no treatment control, 5 μg/mL NNV was the minimum effective concentration able to kill the trypanosome population within 24 hours, whereas 20 μg/mL NNV caused cell death within 8 hours.

**Figure 2.**
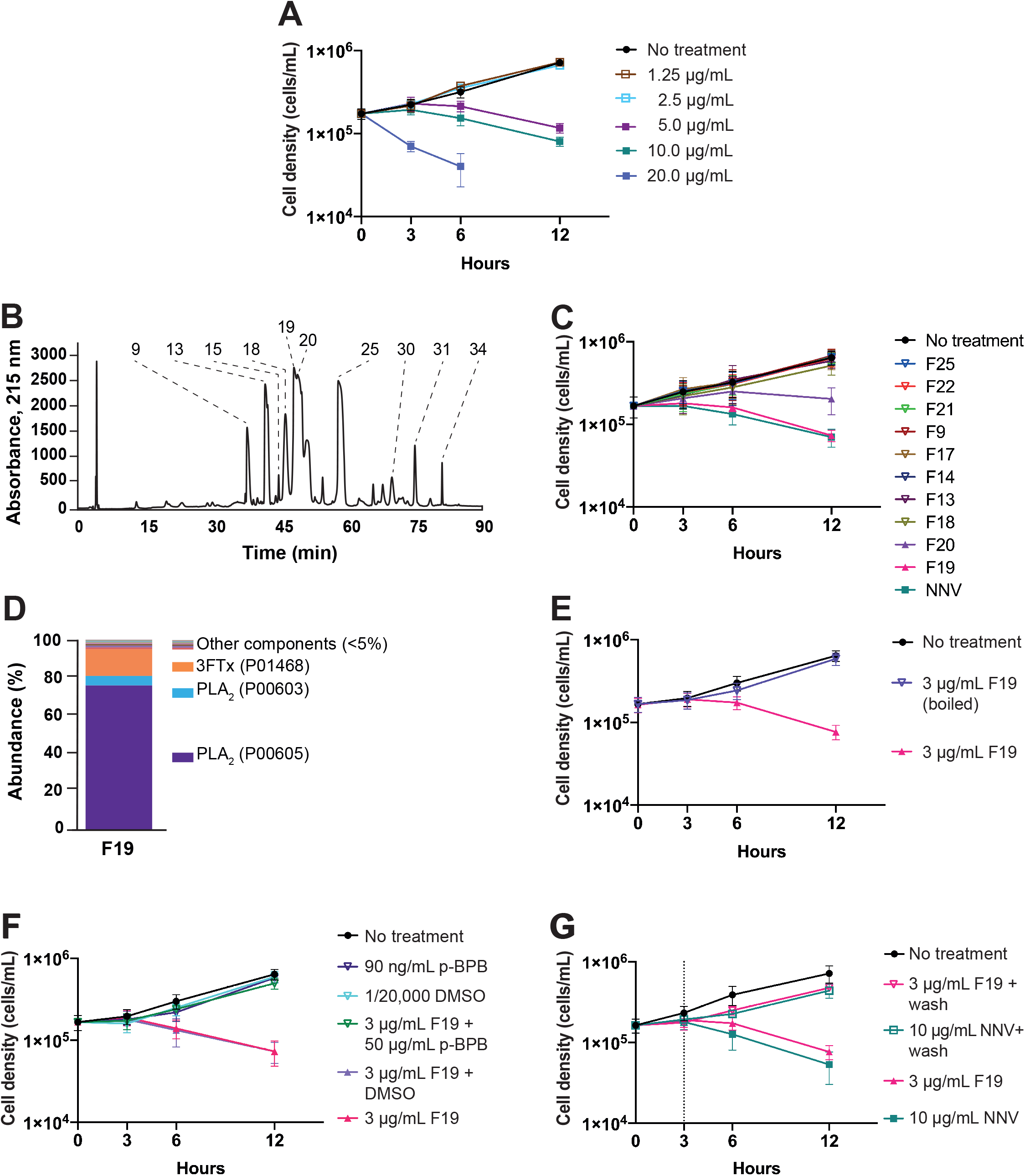
*N. nigricollis* PLA_2_s are the necessary and sufficient venom components to induce trypanocidal activity. A. *N. nigricollis* venom (NNV) causes rapid *T. brucei* death. 20 μg/mL NNV kills the entire population of *T. brucei* within 8 hours. 5 μg/mL NNV was the minimum effective concentration able to kill the cell population within 24 hours. **B.** *N. nigricollis* **venom chromatogram**. NNV was fractionated by RP-HPLC. **C. Fractions containing PLA_2_s retain trypanocidal activity, showing that PLA_2_s are the necessary components to induce cell death.** Fraction 19 containing PLA_2_s and the juxtaposed fraction 20, also containing PLA_2_s, retained trypanocidal activity upon venom fractionation. Venom fractions were tested at 3 μg/mL. NNV was tested at 10 μg/mL. **D. Fractions retaining lethal effects contain PLA_2_s.** LC-MS analysis of fraction 19. The relative abundance of PLA_2_s in fraction 19 is 83%. **E. PLA_2_s require tertiary structure to exert their trypanocidal activity.** 3 μg/mL. Fraction 19 was incubated at 100 °C for 5 minutes and then further incubated for 12 hours with cultures of *T. brucei* VSG121 wild type. Boiled fraction 19 imperiled trypanocidal activity. **F. PLA_2_s require their intact catalytic activity to exert trypanocidal activity.** Fraction 19 was incubated for 1 hour at room temperature with 50 μg/mL *p*-bromophenacyl bromide (*p*-BPB), and further incubated with *T. brucei* cultures. for 12 hours. Chemical modification of the PLA_2_s catalytic His-47 rescued cell growth. **G. PLA_2_ action is sufficient to induce trypanocidal activity.** Cultures of *T. brucei* VSG121 wild type were incubated with 3 μg/mL fraction 19 for 3 hours. Then, cells were washed in fresh HMI-9 media. Finally, cells were resuspended in further 10 mL HMI-9 media and incubated up to 12 hours. Cells recover upon media replacement, confirming the trypanocidal activity is caused by fraction 19 in a time-dependent manner (N = 3).

### *N. nigricollis* PLA_2_s are necessary and sufficient to induce trypanocidal activity

To elucidate which of the venom components were responsible for *T. brucei* cell death, NNV was fractionated by reversed-phase HPLC (Figure 2 B) and individual fractions were tested for trypanocidal activity. Cultures of *T. brucei* were incubated for 24 h with fractions 9, 13, 15, 18, 19, 20, 21, 25, 30, 31, and 34 as these contained the most abundant venom proteins (Fig. 2 C). Growth was compared to no treatment and NNV (10 μg/mL) controls. In one set of growth assays, fractions were tested at 3 μg/mL protein, corresponding to the maximum relative abundance of a known single fraction (Fig. 2 C). Fraction 19 killed trypanosomes with the same potency and in a similar timeframe compared with NNV. Fraction 20 slowed down growth rate but was not as effective as fraction 19. No other fraction affected cell survival or growth rate at the tested concentrations.

Fraction 19 was then characterised by LC-MS to analyse its composition in full (Fig. 2 D). LC-MS analysis found 150 peptide groups from 10,256 peptide spectrum matches, belonging to 33 master protein groups. Out of these, 12 were identified with two or more unique peptides. Label free quantification showed two phospholipase species (UniProt codes: P00605 and P00603), and one cytotoxin (UniProt code P01468) being the most abundant proteins in the fraction. In total, quantification indicated that phospholipase proteoforms accounted for approximately 83% of the protein content in the fraction, followed by cytotoxins at 15%.

We proceeded to investigate the role of PLA_2_s in trypanosome cell death. PLA_2_s were inactivated by two methods. First, fraction 19 was incubated at 100 °C for 5 minutes prior to incubation at 3 μg/mL with cultures of *T. brucei* for 24 hours (Fig. 2 E). The growth was measured and compared to no treatment and native fraction 19 controls. After heat treatment, fraction 19 had no effect on cell proliferation. Second, fraction 19 was incubated with 50 μg/mL *p*-bromophenacyl bromide (*p*-BPB), an inhibitor for PLA_2_s that covalently binds the catalytic histidine 47^19^. (Fig. 2 F). Cell proliferation was measured and compared to a set of controls: i) no treatment; ii) *p*-BPB; iii) DMSO (*p*-BPB solvent); iv) DMSO + fraction 19; and v) fraction 19. When Fraction 19 was incubated with *p-*BPB, it did not interfere with cell proliferation, indicating that the trypanocidal activity of the PLA_2_s was lost through selective inhibition. Third, cultures of *T. brucei* were incubated with fraction 19 for 3 hours (Fig. 2 G). Then, cells were washed in fresh HMI-9 media and resuspended in further 10 mL HMI-9 media. Last, cells were incubated for an extra 5 hours. Cell proliferation was measured and compared to a set of controls: i) no treatment and ii) no-washing in fresh media. Cell proliferation was recovered upon removal of media containing PLA_2_s. These observations indicate that PLA_2_s were the necessary and sufficient NNV component to induce trypanosome cell death.

### PLA_2_s induce an enlargement of the flagellar pocket

Upon confirmation that PLA_2_s cause cell death, changes in cell morphology over a time course after addition of PLA_2_s were investigated. Initial experiments were designed to investigate potential PLA_2_ effects on the cell surface. A *T. brucei* cell line expressing a cell surface GPI-anchored eGFP in a VSG121 background was used to visualise the cell surface coat. Cultures of *T. brucei* BSF VSG121 GFP-GPI were incubated with 20 μg/mL NNV for 6 hours (Fig. 3 A).

**Figure 3.**
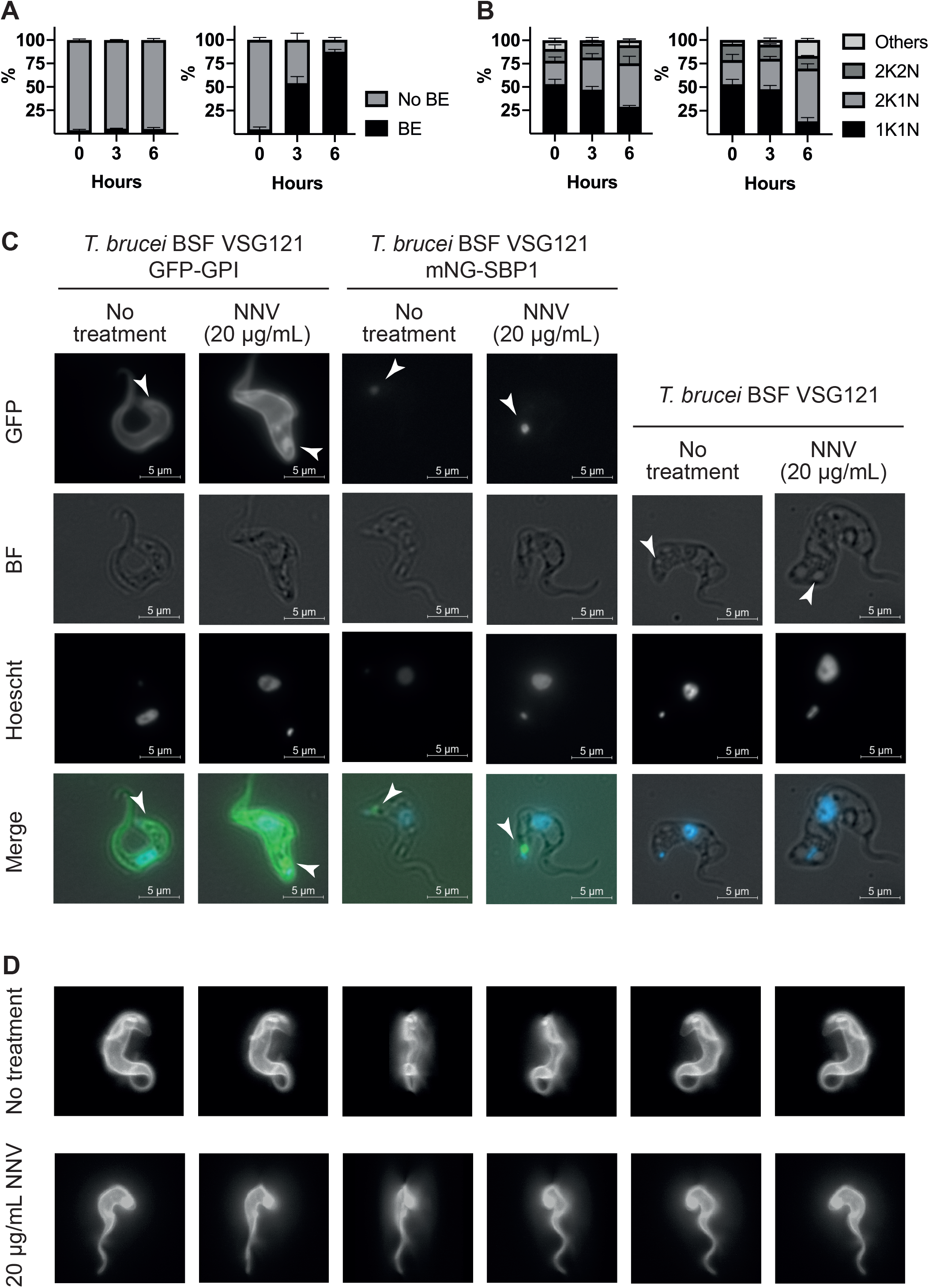
PLA_2_s cause enlargement of the flagellar pocket. A. PLA_2_s cause Big Eye phenotype on *T. brucei*. Cultures of *T. brucei* VSG121 GFP-GPI were incubated with 20 μg/mL NNV, and the absence or presence of Big Eye phenotype (BE) was recorded for 300 cells (N = 3). Left graph, no treatment; right graph, 20 μg/mL NNV. At 3 hours post-NNV addition, over 50% of the cells present BE. 6 hours post-NNV addition, over 85% cells display BE. **B. PLA_2_s compromise cell cycle progression**. Cultures of *T. brucei* VSG121 wild type were incubated with 20 μg/mL NNV. Samples were taken at 0, 3, and 6 hours post-NNV addition. The cell cycle stage (1K1N, 2K1N, 2K2N, other) was annotated for 300 cells (N = 3). Left graph, no treatment, right graph, 20 μg/mL NNV. A 25% increase of 2K1N cells and a 4-fold increase of cells fitting within the ‘other’ category (i.e. cell cycle abnormalities) were observed as a consequence of the PLA_2_ effect. These observations indicate cells try to progress through the cell cycle up to the duplication of the kinetoplast, and then they cannot proceed any further. **C. SBP1-mNG distribution indicated an expansion of the flagellar pocket membrane area flagellar pocket.** Fixed fluorescence microscopy was performed prior to NNV addition and 6 hours post-NNV addition to *T. brucei* BSF VSG121 GFP-GPI, *T. brucei* BSF VSG121 SBP1-mNG, and *T. brucei* BSF VSG121. Accumulation occurs at the flagellar pocket, as it can be observed in the GFP panels. Accumulation is not a by-product of GFP or mNG expression as it can be observed in DIC images for wild type cells as well as in the modified cell lines. **D. Accumulation occurs at the flagellar pocket membrane**, not passively at the empty space created by the flagellar pocket membrane. Fixed confocal fluorescent microscopy was used to create 3D reconstructions of the cells with and without NNV (6 hours post-addition). The panels represent snapshots of the 3D reconstruction twisting along the Y-axis clockwise.

Cell morphology was recorded prior to NNV addition, 3 hours and 6 hours post-NNV addition, and compared to no treatment conditions. At 3 hours post-NNV addition, around 50% of the trypanosomes had an enlarged flagellar pocket similar to the ‘Big Eye’ phenotype caused by blocking endocytosis. At 6 hours, over 80% of the cells displayed the Big Eye phenotype.

In a second set of experiments, the effect of NNV on the cell cycle was analysed (Fig. 3 B). This experiment was carried out in the same conditions as the first, except wild type VSG121 expressing cells were used and Hoechst 33342 was used to visualise the nucleus and kinetoplast. At 6 hours post-NNV addition, a 25% increase in cells with 2 kinetoplasts (2K1N) was observed when compared to no treatment conditions. In addition, a 4-fold increase in cells presenting a range of cell cycle abnormalities was observed, such as incomplete cell division resulting in two daughter cells remaining joined by the posterior ends and cells with only one kinetoplast but two nuclei.

The effect on the flagellar pocket was investigated further by making a cell line with one endogenous allele of syntaxin binding protein 1 (SBP1, Tb927.9.1970) tagged at the C-terminus with mNeon Green fluorescent protein (mNG) as a flagellar pocket membrane marker. Cultures of *T. brucei* BSF VSG121 expressing SBP1-mNG were incubated with 20 μg/mL NNV for 8 hours. Cell morphology was analysed by fluorescence microscopy prior to NNV addition and 6 hours post-NNV addition, and compared to no treatment conditions. *T. brucei* BSF VSG121 GFP-GPI and *T. brucei* BSF VSG121 wild type were treated in parallel (Fig. 3 C). The flagellar pocket in the SBP1-mNG expressing cells was enlarged, indicating that *N. nigricollis* PLA_2_s disrupt normal flagellar pocket function.

To further characterise the nature of the accumulation causing an enlarged flagellar pocket, confocal microscopy was performed on *T. brucei* BSF VSG121 GFP-GPI cells 6 hours after 20 μg/mL NNV addition and compared to no treatment control (Fig. 3 D). The flagellar pocket in NNV-treated cells adopts a sphere-like shape, and the accumulation of GFP-GPI is located at the membrane, as opposed to being released from the membrane and accumulating in the interior space.

### PLA_2_s inhibit endocytosis

The next set of experiments investigated the nature and origin of the flagellar pocket enlargement. First, membrane accumulation could be the result of newly synthesised plasma membrane being exported to the flagellar pocket but not released onto the cell body. Alternatively, enlargement could result from an inhibition of endocytosis. To distinguish between these two possibilities, protein translation was inhibited using cycloheximide (CHX) prior to addition of NNV^20^.

First, cultures of *T. brucei* BSF VSG121 wild type were incubated with 50 μg/mL CHX for 1 hour, and then 20 μg/mL NNV was added. Cells were incubated for a total of 9 hours. Cell growth was measured prior to CHX addition and 1, 4, and 7 hours post CHX addition. A set of controls was carried out in parallel: i) no treatment; ii) 50 μg/mL CHX, and iii) 20 μg/mL NNV. As expected, CHX slowed down the growth rate (Fig. 4 A). The growth rate in the culture incubated with NNV and CHX was less slowed down than in those with only NNV. The implications of this observation are discussed in the following section.

**Figure 4.**
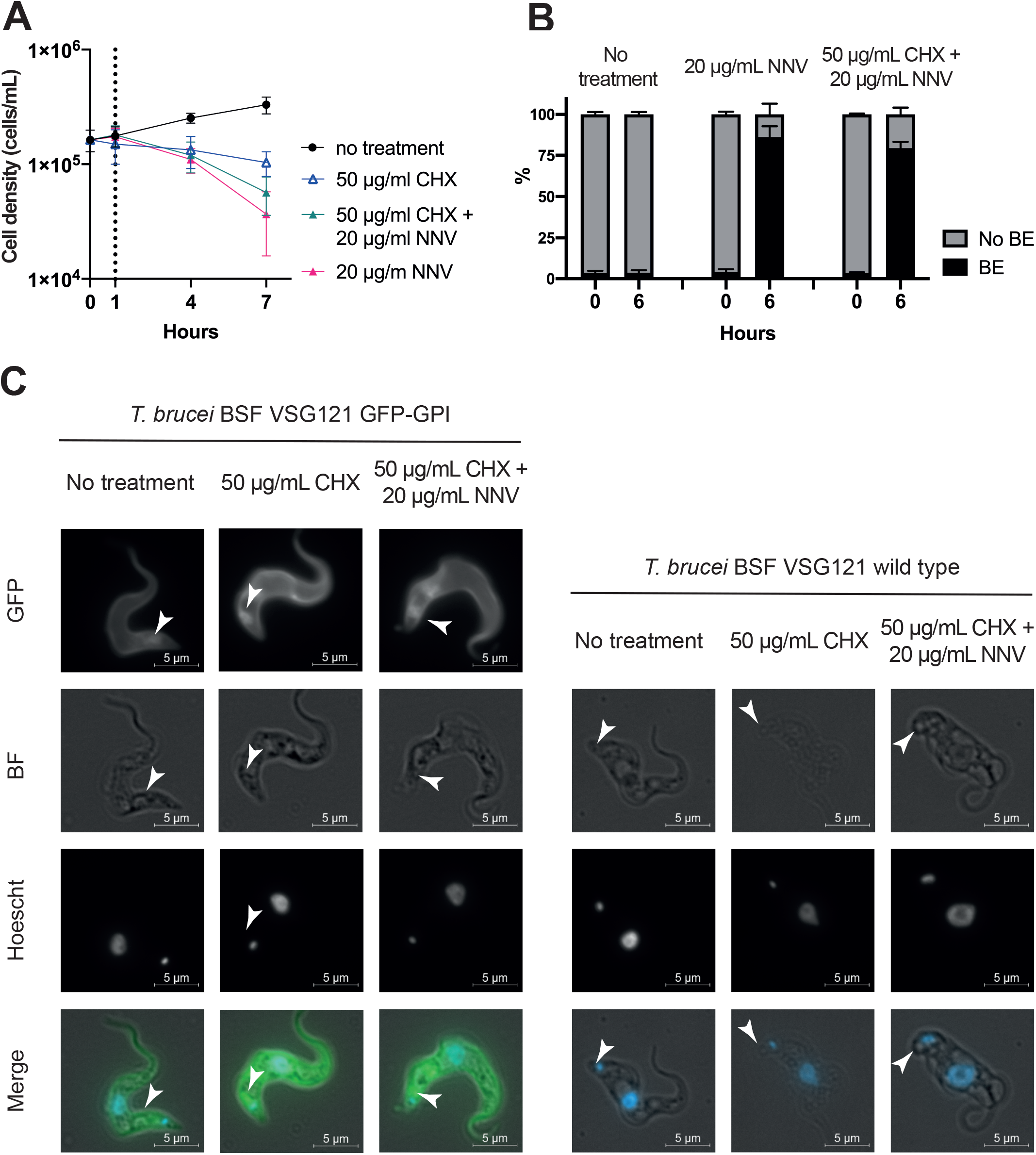
PLA_2_s inhibit endocytosis of *T. brucei* plasma membrane components at the site of the flagellar pocket. A. Protein synthesis inhibition lessens the cell growth phenotype caused by PLA_2_ toxicity. Cells were incubated with cycloheximide (CHX), NNV, or both, and cell growth was monitored. *T. brucei* BSF VSG121 wild type cells incubated with CHX/NNV die more slowly than those cells incubated with NNV only. **B. CHX reduces the proportion of cells presenting Big Eye phenotype (BE) caused by the action of PLA_2_s by 15-25%.** *T. brucei* BSF VSG121 GFP-GPI cells were incubated with identical conditions as in Fig. 4 A, and the absence or presence of BE in 300 cells (N = 3) was annotated utilising live fluorescence microscopy. **C. Blockade of inside-out cargo trafficking relieves BE phenotype intensity.** Fixed fluorescence microscopy was performed at 3 and 6 hours post-NNV addition (equivalent to 4 and 7 hours post CHX addition) on *T. brucei* BSF VSG121 wild type and *T. brucei* BSF VSG121 GFP-GPI. A milder (i.e. smaller) BE phenotype could be observed when cells were treated with CHX/NNV, confirming that the PLA_2_ blockade effect occurs at the endocytosis pathway.

In a second set of experiments, *T. brucei* BSF VSG121 SBP1-mNG cells were incubated with 50 μg/mL CHX for 1 hour, and then 20 μg/mL NNV was added. Cell morphology was analysed at 4 hours and 7 hours after CHX addition. A set of control conditions were carried out in parallel: i) no treatment and ii) 20 μg/mL NNV (Fig. 4 B). The proportion of cells with an enlarged flagellar pocket caused by NNV was similar with and without CHX. This observation implied that flagellar pocket enlargement more likely occurred due to inhibition of endocytosis.

*T. brucei* BSF VSG121 wild type and *T. brucei* BSF VSG121 GFP-GPI treated as above were imaged by fluorescence microscopy (Fig. 4 C). As described before, regardless of protein synthesis inhibition, the enlarged flagellar pocket persisted throughout the tested cell lines. No morphological effect was detected on cells treated with CHX only, but trypanosomes died within 24 hours.

### The flagellar pocket is the primary target for toxic effect of PLA_2_s

In the experiments above, PLA_2_ treatment seemed to cause a reduced density of GFP-GPI on the plasma membrane covering the cell body. We investigated whether this decrease represents a loss of GPI-anchored proteins, possibly due to PLA_2_ action (Fig. 5 A), or a redistribution to the flagellar pocket. Cultures of *T. brucei* BSF VSG121 wild type, VSG121 GFP-GPI, and VSG121 SBP1-mNG were incubated with 20 μg/mL NNV with or without one hour pretreatment with CHX and samples were analysed by flow cytometry prior to NNV addition and 6 hours post-NNV addition (Fig. 5 B). The total cell fluorescence was unchanged, except for a slight decrease in the double treatment group. This observation indicates there is a redistribution of cell surface proteins to the flagellar pocket rather than a loss.

**Figure 5.**
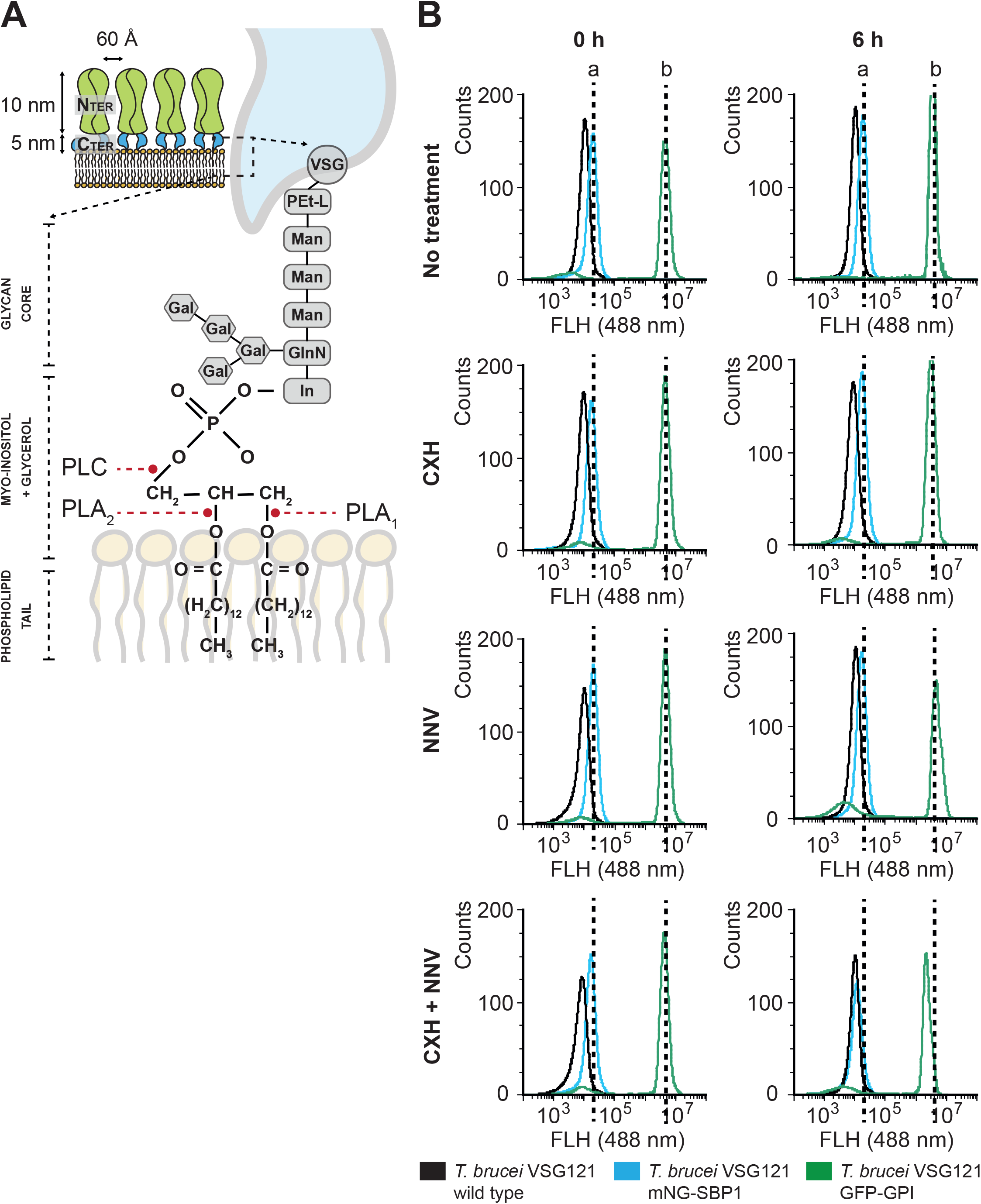
PLA_2_ action is not sufficient to release GPI-anchored proteins. A. schematic representation of the GPI-anchor linking VSG C-terminus to the plasma membrane. The GPI anchor binds an aspartic acid from the VSG structure by a polyethylene glycol linker. This connects to a glycan core (three mannoses and one glucosamine branch linked to four galactoses). The glucosamine is bound to myo-inositol, which connects to the phospholipid tails. The 2-acyl bond is the canonical target for PLA_2_s, whereas PLA_1_s hydrolyse 1n-acyl bonds. *T. brucei* PLC is known to hydrolyse phosphate bonds linking myo-inositol to glycerol. **B.** Live flow cytometry on *T. brucei* BSF VSG121 wild type (black), *T. brucei* BSF VSG121 mNG-SBP1 (blue), and *T. brucei* BSF VSG121 GFP-GPI (green). *T. brucei* BSF VSG121 wild type display 10^3^ FAU due to its own autofluorescence. NNV (i.e. PLA_2_s) does not lead to a loss of fluorescence by hydrolysing GFP-GPI from the cell surface. In contrast, it allows a redistribution effect: Due to the hydrodynamic forces originating from the cell swimming, GPF-GPI is relocated to the flagellar pocket. There, endocytosis is blocked due to the PLA_2_ blockade effect and, thus, GFP-GPI accumulates. There is not a significant loss of fluorescence, but a relocation from the cell surface to the flagellar pocket. Blocking protein synthesis (CHX) in combination to NNV leads to a reduction in the total fluorescence intensity as i) there is no GFP-GPI being transported to the cell surface, and ii) the GPF-GPI in the cell surface is being relocated to the flagellar pocket. In summary, these observations underpin the hypothesis that trypanocidal effect is caused by endocytosis ablation by PLA_2_ action, enhanced by protein redistribution to the flagellar pocket, and not GPI hydrolysis.

### PLA_2_s compromise Rab11-mediated recycling pathway fitness

To evaluate whether other subcellular compartments involved in endo- and exocytosis are affected by NNV, a set of cell lines with different mNG-tagged subcellular compartments were used. Rab5A (*T. brucei* BSF VSG121 mNG-Rab5A) and Rab5B (*T. brucei* BSF VSG121 mNG-Rab5B) were tagged for the early endosome, Rab7 (*T. brucei* BSF VSG121 mNG-Rab7) for the late endosome, and Rab11 (*T. brucei* BSF VSG121 mNG-Rab5A) for the recycling endosome.

Cultures of these *T. brucei* cell lines were incubated with 20 μg/mL NNV and imaged by fluorescence microscopy 6 hours after NNV addition (Fig. 6). No observable effect was found in the early endosome (Rab5A and Rab5B) or late endosome (Rab7). In contrast, a slight distortion of the Rab11 wild type pattern was caused by NNV. Rab11 appeared in a more dispersed pattern, as opposed to the no treatment control, where Rab11 was located in discrete spots. This observation indicates that the recycling route is affected by PLA_2_ activity.

**Figure 6.**
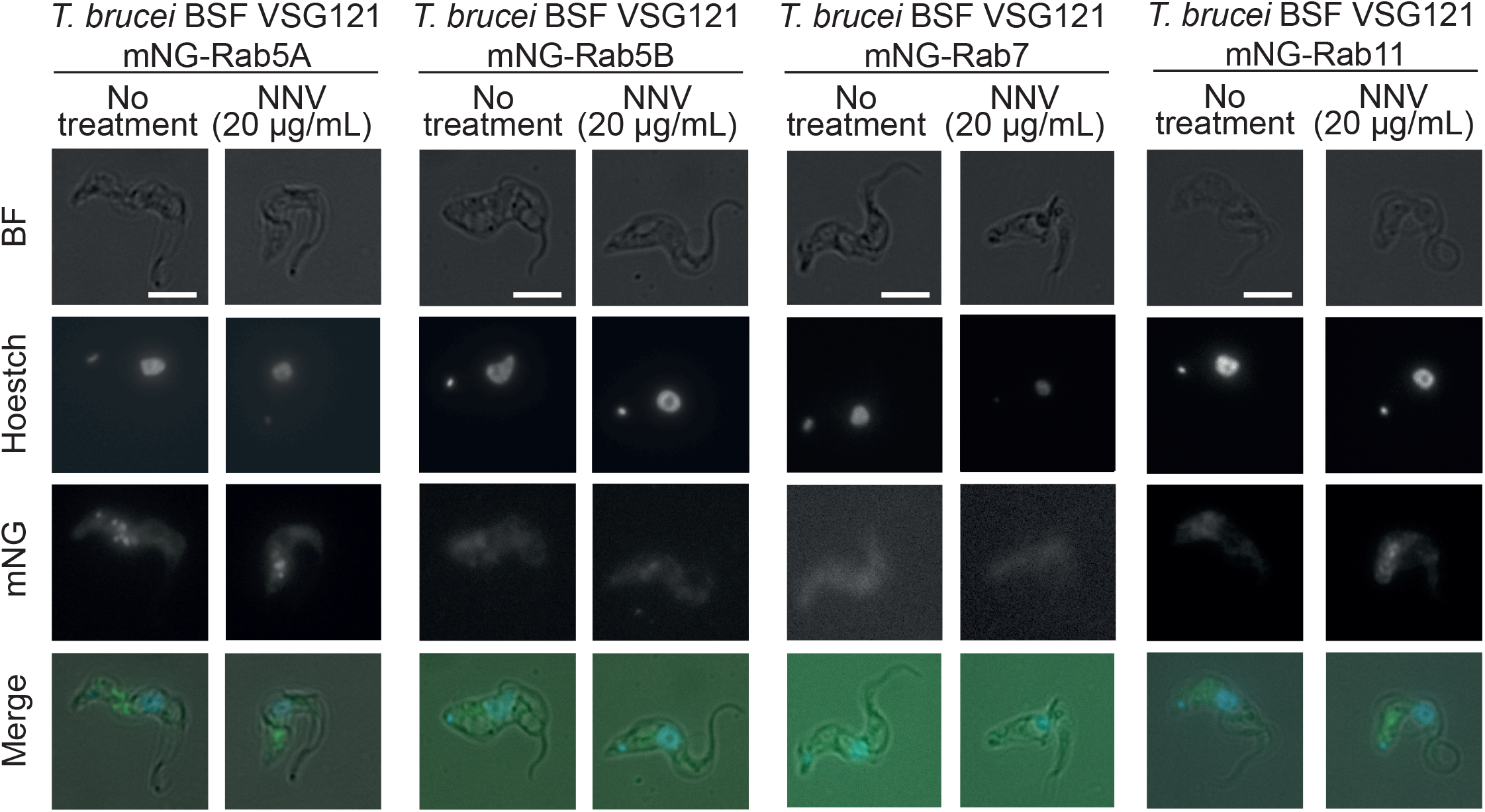
The downstream Rab11+ recycling endosome is affected by PLA_2_ action. Fixed fluorescence microscopy (100X) was performed at 6 hours post-NNV addition and no treatment conditions. 100 cells were observed per experiment (N=3). Rab5A+/5B+ and Rab7+ compartments remain unaffected, whereas the Rab11+ recycling endosome displayed a discrete and bright spot pattern as opposed to a more disperse one (no treatment). This provides an indication of the downstream consequences of the PLA_2_ effects. Due to cargo accumulation at the flagellar pocket, the recycling endosome struggles to deliver recycled cargo back to the flagellar pocket, and its morphology is observably altered. Scale bar = 5 μm.

## Discussion

In the mammalian bloodstream, the trypanosome cell surface is covered by a densely packed coat of VSG molecules. As individual bloodstream form trypanosomes grow, newly synthesised VSG-coated plasma membrane is added to the existing plasma membrane. At the same time, pre-existing VSG-coated plasma membrane is endocytosed from the flagellar pocket and returned after a circular digression around the endosomal system. The processes of biogenesis, exocytosis, endocytosis, and recycling of VSG molecules are pivotal to trypanosome survival^21^, as VSG represents over 10% of the total protein content and more than 90% of surface proteins. Remarkably, trypanosomes have the ability to recycle the entire surface coat within 12 minutes^7^. The flux of recycling VSG is at least 10 times greater that the flux of newly synthesised VSG reaching the cell surface^7^. Here, we tested the effect of the hydrolytic enzymes in snake venom on the VSG coat.

Snake venoms are heterogeneous cocktails of proteins, including different snake venom metalloproteinases (SVMPs), snake venom serine proteinases (SVSPs), phospholipases A_2_ (PLA_2_s), three-finger toxins (3FTxs), and other minor components^22^. In particular, *Naja nigricollis* venom (NNV) was chosen as: (i) It is very active against cellular membranes in the eyes, the main target of venom being spat at a predator threatening a spitting cobra^23^. (ii) Cobras, such *N. nigricollis*, *N. katiensis*, *N. pallida,* and *N. nubiae,* have a significant proportion of PLA_2_s in their venom compositions (20 to 30%). (iii) Unlike many other venoms, NNV composition has been characterised^17^, and (iv) *N. nigricollis* PLA_2_s have been structurally^15^ and functionally^24^ characterised. In this study, we observed that the effects of the PLA_2_-enriched fraction 19 from NNV on trypanosomes can be blocked by the inhibitor *p-*bromophenacyl bromide, which has been extensively used to covalently modify the catalytic His-47 in PLA_2_s and ultimately alter their tertiary structure^25^. This provides strong evidence that PLA_2_s are the NNV components causing flagellar pocket enlargement and subsequent cell death.

Snake venom PLA_2_s have evolved to attack plasma membrane components, and it was originally expected that a hydrolytic effect would have been observed on the GPI-anchor of VSG. However, the VSG, or at least GPI-anchored GFP, was unaffected by addition of exogenous PLA_2_s, and instead a phenotype associated with disruption of endocytosis was observed. The observation that the effects of the PLA_2_-enriched fraction 19 on trypanosomes can be blocked by the inhibitor *p-*bromophenacyl bromide further provided strong evidence that PLA_2_s are the NNV components causing flagellar pocket enlargement and subsequent cell death. The mechanistic cause of the enlargement of the flagellar pocket remains unclear. The enlargement is not dependent on new protein synthesis, and this suggests a block in endocytosis, as reported for RNAi suppression of the clathrin light chain^8^ or actin^18^. There was little effect on internal membrane compartments as judged by tagging various markers of internal compartments of the endosomal pathway. The exception was a change in the distribution of Rab11, which marks recycling vesicles^9^. This could be secondary as, if no endocytosis is occurring, there is nothing to recycle.

The cell cycle was disrupted by NNV addition. There was an abnormal proportion of cells showing two kinetoplasts and a two-fold increase in cells with two flagellar pockets. These observations suggest an arrest in, or cells taking longer time than usual to traverse, the latter stages of the cell cycle. RNA interference of VSG mRNA causes an arrest in the early stages of cytokinesis possibly due to a shortage of new VSG-coated plasma membrane^26^, and the effect of NNV may also reflect a plasma membrane defect. In cells with two flagellar pockets, only one of the pockets is enlarged by NNV, possibly reflecting a lack of activity on the new pocket.

Venoms have been used in the past as molecular tools for the study of trypanosomatid-related diseases^11–13^. However, the mechanisms leading to trypanocidal effects have never been characterised. Our work constitutes the first morphological and mechanistic description of the toxicity triggered by *N. nigricollis* PLA_2_s in *T. brucei*. We provide a description of a novel mechanism by which trypanosomes die. *N. nigricollis* PLA_2_s cause cargo accumulation at the flagellar pocket, thus blocking endocytosis, a high-rate constitutive process for the cell, which is essential for parasite survival through immune evasion. Targeting trafficking pathways has been suggested as a potential alternative to improve treatment of Human African Trypanosomiasis^27^. While we remain skeptical that snake PLA_2_s could serve as novel drug leads, the data presented in this study shine light on novel mechanisms for targeting and killing trypanosomes, which could potentially be exploited therapeutically.

## Materials and methods

### Snake venom

*Naja nigricollis* venom (NNV) from Tanzanian specimens was purchased in lyophilised form from Latoxan, France (cn. L1327B). Venom was reconstituted in HMI-9 media without fetal calf serum (FCS) when used in trypanosome cell culture.

### Venom fractionation and proteomics characterisation

Whole venom fractionation was performed by reversed-phase high performance liquid chromatography (RP-HPLC) (Agilent 1200) on a C18 column (250 × 4.6 mm, 5 μm particle; Teknokroma), using HPLC Chromeleon software^28^. 2 mg/mL whole venom in phosphate buffered saline (PBS: 137 mM NaCl, 3 mM KCl, 8 mM Na_2_HPO_4_.2H_2_O, 1.4 mM KH_2_PO_4_) was loaded onto the column and run through an acetonitrile/trifluoracetic acid gradient for elution. Then solvent was evaporated using a speedvac. Finally, equivalent fractions from different rounds were pooled together in HMI-9 media without foetal calf serum (FCS). Fraction were analysed by SDS-PAGE (7%) and silver stained^29^.

20 μg of fraction 19 were reduced with 5 mM Tris(2-carboxyethyl)phosphine (646547, Sigma Aldrich) for 60 min at 65 °C, and afterwards alkylated with 20 mM 2-chloroacetamide (C0267, Sigma Aldrich) for 30 min at 65 °C. Proteins were digested with trypsin (V5280, Promega) overnight, with a trypsin:protein ratio of 1:50. Desalting was performed with in-house packed C18 disk tips following standard reversed phase SPE protocols for sample cleanup^30,31^. LC-MS analysis was performed with 300 ng of sample injected in a Q Exactive hybrid quadrupole-Orbitrap mass spectrometer (Thermo Scientific), using an EASY-nLC 1200 System (LC140, ThermoFisher Scientific), and an EASY-Spray column (50 cm × 75 μm ID, PepMap RSLC C18, 2 μm ES803, Thermo Scientific), with a 70 min gradient length of increasing acetonitrile percentage. The mass spectrometer was operated in data dependent acquisition mode, scan range of 300-1750 m/z with MS resolution of 70,000, AGC target of 3e6 and maximum injection time of 20 ms. MS2 scans were obtained for top 10 precursors, with first mass of 120 m/z, resolution of 17,500, AGC target of 1×10^6^, maximum injection time of 60 ms, 1.6 m/z isolation window, and NCE of 25.

Data analysis was conducted with Proteome Discoverer 2.4 (Thermo Scientific), with workflows adjusted for label free quantification. The FASTA database used was extracted from the reviewed snake venom protein list from the UniProt database (taxonomy:“Serpentes (snakes) [8570]” annotation:(type:“tissue specificity” venom) AND reviewed:yes) containing 2,330 venom proteins. Full digestion with trypsin was indicated in the settings, with two maximum missed cleavages allowed, and peptide length of 6-144. Precursor mass tolerance and fragment mass tolerance were set to 10 ppm and 0.02 Da, respectively. Oxidation (methionine) and deamidation (glutamine and asparagine), and acetylation (protein N-terminus) were added as variable modifications, while carbamidomethylation (cysteine) was added as static modification. Summed peptide area abundances were used to calculate master protein abundance (i.e. the protein chosen to represent the group in the cases where peptides identified match to multiple proteins), with filtering only for high confidence peptides (FDR 0.01). Data interpretation, processing and plot generation were performed in Python 3.6 programming language.

### Trypanosome cell culture

*T. brucei* Lister 427 VSG121 BSF cells were maintained in culture at a density of 1 × 10^5^ to 2 × 10^6^ cells/mL in HMI-9 salts supplemented with 10% (v/v) heat-inactivated fetal calf serum, penicillin (100 U/mL), and streptomycin (10 μg/mL), and kept at 37 °C and 5% CO_2_. Cell counts were carried out using a hemocytometer.

### Cell line engineering

A strategy using long oligonucleotide primers was used to tag endocytic pathway components^32^. In brief, forward primers contained the last 80 base pairs of the 5’ UTR of the target gene and the 20 base pairs of the 5’ pPOTv7 primer binding sequence; reverse primers contained the 20 base pairs of the 3’ pPOTv7 primer binding sequence and the first 80 base pairs of the target gene including the start codon, in the reverse complement^32^. PCR amplification from the pPOTv7 plasmid used resulted in a product carrying blasticidin resistance and mNEON green (mNG) protein. The 5’ UTR sequences were obtained from the TREU 927 reference genome for Tb927.10.12960 (Rab5A), Tb927.11.4570 (Rab5B), Tb927.8.4620 (Rab7), and Tb927.8.4330 (Rab11). PCR products were introduced into *T. brucei* Lister 427 expressing VSG221 by standard procedure and transformants were selected with 10 μg/mL of blasticidin. Expected localisation of N-terminally tagged endocytic pathway proteins was confirmed by fluorescence microscopy. Following the same method, *T. brucei* BSF Lister 427 cell line expressing VSG121 mNG-SBP1 was developed by C-terminally tagging SBP1 (Tb927.9.1970) with mNG. Successful tagging was confirmed by fluorescence microscopy.

*T. brucei* Lister 427 cell line expressing VSG121 GFP-GPI was engineered by introducing a transgene into the VSG121 expression site using the same approach as used to insert a second VSG^33^ The plasmid p4716 was cut with Acc65I and SacI to release the insert which was then used for transfections. The sequence of the insert is available in Fig. S1.

### Cell morphology analysis

The morphology of all cell lines was analysed by fluorescence microscopy using Zeiss Axioimager M1. Images were processed using Zeiss AxioVision v5.6.1, analysed by NIH Image J, and made into figures in Adobe Illustrator.

For live cell microscopy, cells were incubated with 20 μg/mL of NNV and 1 mL of cells was harvested at 3 and 6 hours post-NNV addition. The sample was centrifuged for 1 minute at 3,000 rpm in an Eppendorf microfuge, the supernatant was removed, and the pellet was resuspended in 10 μL HMI-9. 4 μL were pipetted onto the microscope slide and covered by a 25 × 50 mm coverslip for visualisation. Untreated cell cultures were analysed alongside as negative control. For fixed cell microscopy, the procedure was identical, but 1% formaldehyde was added to the 10 μL resuspended cell pellet prior preparing the slide. Confocal microscopy was performed on live cells using following the same method described above. Both the 2-dimensional images and the Z-stacks were processed by NIH Image J.

### Cell cycle analysis

1 mL of cells was harvested from the trypanosome culture and recovered by centrifugation at 10,000 rpm for 1 minute. The supernatant was removed, 10 μL of HMI-9 was added, and the pellet was resuspended. 4% (w/v) paraformaldehyde was added to fix the cells. Finally, 1/100,000 Hoechst 33342 was added. Fluorescent microscopy was then performed to visualise kinetoplast and nuclei in the cells.

### Cell surface analysis

Cell surface fluorescence levels due to GFP-GPI were analysed by flow cytometry using a CytoFlex Flow Cytometer (Dept. of Pathology, University of Cambridge). *T. brucei* Lister 427 VSG121 (wild type) and *T. brucei* Lister 427 VSG121 p4716 (GFP-GPI) cells were incubated with 20 μg/mL NNV, and 1 mL of cells was taken at 3 and 6 hours post-NNV addition and transferred to a capped glass tube. Cells not subjected to NNV were also analysed in parallel. Wild type cells were used as a negative control for GFP fluorescence. The cytometer was set to record 10,000 events at 60 μL/min sample intake. A blank sample (pure water) was run between samples to avoid cell carry overs. The gating strategy employed aimed at separating single live cells from those dividing, forming clumps while dying, and debris. Data analysis was done using FlowJo software and image processing was done in Adobe Illustrator.

### SDS-PAGE and immunoblotting

Cell lysates were made by harvesting BSF cell culture. After centrifugation (3,000 rpm, 10 minutes), the pellet was resuspended in 1 mL HMI-9 without serum and transferred into a 1.5 mL tube. The sample was centrifuged (10,000 rpm, 1 minute), the supernatant was pipetted out. This step was repeated twice. Finally, the pellet was resuspended in HMI-9 + 1X SDS-PAGE sample buffer to a concentration of 10^8^ cell equivalents/mL and incubated at 95 °C for 5 minutes. Then, SDS-PAGE was carried out^34^. For each sample, 10 μL of cell lysate (2 × 10^6^ cell equivalents/well) was used. Gels were stained with Coomassie, silver staining^29^, or transferred to a membrane. Immunoblotting was carried^35^. Gel analysis was performed by NIH Image J, and the image was processed by Adobe Photoshop.

### *In vitro* trypanosome cell killing assays

All cell lines were incubated with 20 μg/mL *N. nigricollis* whole venom (NNV) or with the relative concentration of each individual venom fraction. Trypanosome cell concentration at the start of the experiment was 1.5 × 10^5^ cells/mL. Growth was measured by counting cells on a hemocytometer. Negative controls were carried out for all cell lines under conditions not containing NNV. The assays were performed in triplicates and collected data were Log_10_ transformed.

### Protein synthesis inhibition assay

*T. brucei* Lister 427 VSG121 (wild type) and *T. brucei* Lister 427 VSG121 GFP-GPI cell lines were incubated with i) 50 μg/mL cycloheximide (CHX); ii) 20 μg/mL NNV; iii) 50 μg/mL CHX + 20 μg/mL NNV; or iv) HMI-9 media only. NNV was added one hour post CHX addition. Trypanosome cell concentration at the start of the experiments was 1.5 × 10^5^ cells/mL. Growth was measured by counting cells on a haemocytometer at 1, 4, and 7 hours post CHX addition. In addition, cell surface fluorescence was analysed by flow cytometry, and total GFP within the cell was analysed by anti-GFP western blot.

## Acknowledgements

We thank Dr. Sam Dean for kindly providing the plasmid pPOTv7. MC is a Wellcome Investigator (217138/Z/19/Z). We thank Wellcome Trust for the funding for this study.

## Contribution

AME, MC, and AHL conceived the study. AME, AHL, OJSM, IM, KK and JAJ performed and analysed the experiments. AME, MC, and AHL wrote the manuscript. All authors contributed to editing the manuscript.

