## Supplemental Figure 1 for "Black-necked spitting cobra (*Naja nigricollis)* phospholipases A_2_ cause *Trypanosoma brucei* death by blocking endocytosis through the flagellar pocket"

# A

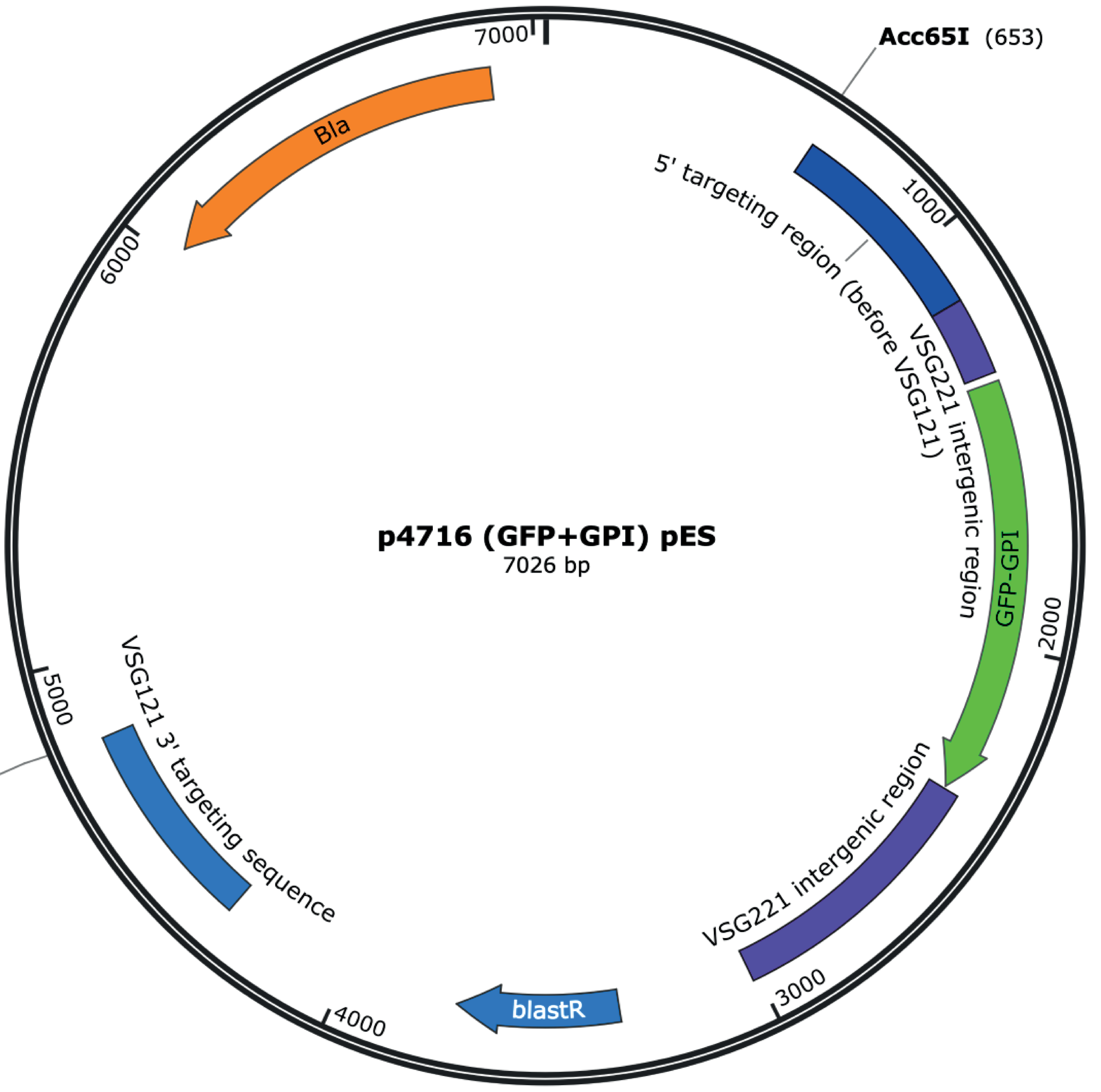


# B

**Acc65I**

**GGTACC**CTGTTCTAAACATAAACATAGCAATAAATATTATCGGAAACGAGCAAATACAGGAAGGTTGTAATTGAAATAGAGTCACTAAATTATCCGGTAAACCAGTGGAAGTAATAGTAAAA
TTAACAAAAGTATAGAATGAAATAATGTGACTGTCAACAACTTCCGCAGAACAAGGGAAAATAATACATGAAGAGGAAGAGGAGATAATGCACCGTATTTCACTCAGAAATGGATATATTA
TTTTGCCGTGGTGGTGGCGTCTATGTTTCGTTGGCTGATGACGGATTCTAATGTTCAGTTGTTGACAATAAGCAACTCCAAAGCACACAATGGTAAACAAAGGAAATGGAACAAAAGG
AAAGTGCTTGAAAAGATTGCAGTCCTACAACAAAATACAAAAATTGCAGTAAGAAAACACAGAGCAGCTGTGATCACTGCCTTGACTGAGGATATATTTTTCGGAATCGAAAAACTAATC
GTTTATAAAATTAGAAAAGCACTGACTCGAGCACACAAGCATTCTATACGTAAAAGATCTAGTATATAGGAGCAACGCTCTGCCAAAACATAATGGCAAGACAAACGGCCGTGTTTGCC
GCTGATGCTACAGAACCAGCTTAATTTCCAGAAGACGAAAATTTGCATGTTTTCCCACAATATTTTAATTACTCTTGAAGATTGTAGTTATTCCTACGCGACACGAACGCGGCATGCCAA
GTAACCAGGAGGCACGTCTTTTTCTTGCAGTGCTTGTGCTTGCACAGGTGCTTCCAATTCTTGTGGATAGTGCAGCAGAGAAGGGTGCACGTGAGGTGCACACAAACCAGGACCCA
AGTGGTAGTAACACAACAAGTGGTAGTAGTTCCATGGGTCGTAAGGGCGAGGAGCTGTTCACCGGGGTGGTGCCCATCCTGGTCGAGCTGGACGGCGACGTAAACGGCCACAAGT
TCAGCGTGCGTGGCGAGGGCGAGGGCGATGCCACCAACGGCAAGCTGACCCTGAAGTTCATCTGCACCACCGGCAAGCTGCCCGTGCCCTGGCCCACCCTCGTGACCACCCTG
ACCTACGGCGTGCAGTGCTTCGCACGCTACCCCGACCACATGAAGCAGCACGACTTCTTCAAGTCCGCCATGCCCGAAGGCTACGTCCAGGAGCGCACCATCAGTTTCAAGGACG
ACGGCACATACAAGACCCGCGCCGAGGTGAAGTTCGAGGGCGACACCCTGGTGAACCGCATCGAGCTGAAGGGCATCGACTTCAAGGAGGACGGCAACATCCTGGGGCACAAGC
TGGAGTACAACTTTAACAGCCACAACGTCTATATCACAGCCGACAAGCAGAAGAACGGCATCAAGGCAAACTTCAAGATCCGCCACAACGTGGAGGACGGCAGCGTGCAGCTCGCC
GACCACTACCAGCAGAACACCCCCATCGGCGACGGCCCCGTGCTGCTGCCCGACAACCACTACCTGAGCACCCAGTCCGTGCTGAGCAAAGACCCCAACGAGAAGCGCGATCAC
ATGGTCCTGCTGGAGTTCGTGACCGCCGCCGGGATCACTCACGGCATGGACGAGCTGTACAAGGGCGCCAAGAAGGTGGCAGATGAGACAGCAAAGGATGGTAAGACAGGTAAC
ACAAACACAACAGGTAGTAGTAACAGTTTTGTGATTAGTAAGACACCACTTTGGCTTGCAGTGCTTCTTTTTTAATTAATTTCCCCCCTCAAATTTCCCCCCTCCTTTTAAAATTTTCCTT
GCTACTTGAAAACTTTTTGATATATTTTAACACCAAAACCAGCCGAGATTTTGTGTTCTGTGTTTTGTAAGTTGACTGTCTGATTGTCTAGAAATATTTTCTGGCAACTAAAATTTTTTTCT
TTTTTCCTGTTTTTTTTGTAGGTAGGTAGGAATGGGGGGGGGGGGTAGTTAGGTAGGTTAGTTAGGTTAGTTAGGGGGGTTAGTTAGGGGGGTTAGGCTTAGGATTAGGATTAGACTTAG
GCTTAGGATTAGGATTAGGATTAGGATTAGGGTTAATTTTTTCCTCTTTTTTTTTAACTCACACCTCTATCCTGGATTTTTAATTTTTTTTTTTTAGCCATTCGCGGCTCCTTTTTTTTTTTTT
GCGCCAATGTTTAATTTTTTATTGTGTTTTCAATTTTTTTGTCAACCATGCAGCGGCTGTTTTGTTATGCGGACCCTAACCCTCCTCCCCCCCCCCCGCCCGCGCACCTCCATTTTTAA
AAATTTTTTTACCGCGTCCTTCAACCAGAATTTTTTTAAATTTTTTTAATTTTTTTTTATTTTCCGTGGTTTTGAATCTTAATTTTTCGGGGGAATTCCTGCAGCCCGGGTAGAAAGTGTGACA
ACGTCGCACCATGTGTAGGTTTTCATTTATGTTCTTTCTTTCTTTTTTTTTGTGAATTTGTTTTCTGTCTCAAATGTTTTTAATTCGCTTGGGACCTATGTTTTTCTTGTTTTTTTGCTCACC
CTTTGTGTAGGAGGCACCCTGTCACGTCTGTGGTTGCGTGTATGCCTTCCTTCCCCTTATTCGCTTCTTCCTGTCGTGTCACACCTCTTTCTCTCTCTCCCTTTCCGCCTTTTCTTTCA
ATCTTGTTTTCTCGACCAGCCCTACTAGAGGAGAAAGAACAGTAACCCTTTCATCAAAGAAAATAGTTCAAACGAATTCATATGCCTTTGTCTCAAGAAGAATCCACCCTCATTGAAAGA
GCAACGGCTACAATCAACAGCATCCCCATCTCTGAAGACTACAGCGTCGCCAGCGCAGCTCTCTCTAGCGACGGCCGCATCTTCACTGGTGTCAATGTATATCATTTTACTGGGGGAC
CTTGTGCAGAACTCGTGGTGCTGGGCACTGCTGCTGCTGCGGCAGCTGGCAACCTGACTTGTATCGTCGCGATCGGAAATGAGAACAGGGGCATCTTGAGCCCCTGCGGACGGTG
CCGACAGGTGCTTCTCGATCTGCATCCTGGGATCAAGGCCATAGTGAAGGACAGTGATGGACAGCCGACGGCAGTTGGGATTCGTGAATTGCTGCCCTCTGGTTATGTGTGGGAGG
GCTAATTCGAATAGACGCGGACGGGGCATTTCCCGTTCGTCATTAGCAGTAGGTAATGAAGATGTTTGTTTCTCGTCCCCTTTCTCCTTCGTCCTTCTGTCATTTTGTTCTTTTGTGTTT
ATGTTTTGTTGTTGTTTTCTTTAATTTTTTTTTTTTCTTCCACGTTTGTGTACATCCGCGCGCCACTCTATTCAGAGAGCCACGGATAGTAGAGGAGGTGGGAAGGGTATATGAGGGACAC
GCGTACCATGATGTGGGATGTATTGGGGTCCCTGTCTGTCCTTACGTGACTATGTATGAACCGTCACGTGTAAGATGAGCTAGTGAGATCAACAGTACAACTCATCAACACGCCTTCTT
CTCGTTAAATGTACACAATCTTGATCCTCCACCTTTATGGGTCCCATTGTTTGCCTCTTCCGCTGTGTGGAGTGCGCCTACACGCACTTCTCACTTCGTAAGTGGTGGTGGCGTAAGTA
TTGCCTAATGTTGACTCTATATTCTCCTCTCCTCACCCCCTCGCGGTGCTGATTTCTGACAGATCTTCAAACACTAGATTAAGCAAAGGACTATTCATCCGTTACTAGTACCACCTGTGC
GACGAAGCTGCAAAAGAATAAAGCGGCACCAAGTAGTTCTAACAAGCTGTAGTGAGACCCAGCTGCACGATAGATTAATTTATTATTTTTTAAATTTTGAGTTTTTTTAAATTTATATAAGT
GTAATTCACCACATTAAAAAGGGGGAAAGAGGACTCAAAATGATATTCTAATTAGCTGAAGGAAACGGTGATGAAAATAGATAATAAACATTCCCCAAAAATATTACCACAACTAACTTCT
TTGTTTTTCTCTTTCATGTTGCTAAACTAGACAACAGCGTTAAGCGATGGCCGTGCACAGAGCCCTAGCGGCGTACGCGATTAGTCTTTACGTTTTACTACCCAGAAAATCGGGAGCA
ACAGACAAAGGCGCGATCAAGTTTGAGACGTGGGAGCCGCTCTGTTTACTGACACAAGACTTCGGTAACCTTTACAACAGAGCGCACAAACTTAATCTCGACATCGACACCTACGTA
ACCGCAGCCCACCGCGGTG**GAGCTC**

**SacI**

**Supplementary Figure 1. A. p4716 (GFP-GPI) pES plasmid map. B. Acc65I/SacI insert used for transfection.**
